# Improved state-level influenza activity nowcasting in the United States leveraging Internet-based data sources and network approaches via ARGONet

**DOI:** 10.1101/344580

**Authors:** Fred S. Lu, Mohammad W. Hattab, Leonardo Clemente, Mauricio Santillana

**Affiliations:** Computational Health Informatics Program, Boston Children’s Hospital,, Boston, MA, USA; Wyss Institute for Biologically Inspired Engineering, Harvard Medical School, Boston, MA, USA; Tecnológico de Monterrey, Monterrey, Mexico; Department of Pediatrics, Harvard Medical School, Boston, MA, USA

**Author notes:** Corresponding author: Mauricio Santillana.

## Abstract

In the presence of population-level health threats, precision public health approaches seek to provide the right intervention to the right population at the right time. Accurate real-time surveillance methodologies that can estimate infectious disease activity ahead of official healthcare-based reports, in relevant spatial resolutions, are critical to eventually achieve this goal. We introduce a novel methodological framework for this task which dynamically combines two distinct flu tracking techniques, using ensemble machine learning approaches, to achieve improved flu activity estimates at the state level in the US. The two predictive techniques behind the proposed ensemble methodology, named ARGONet, utilize (1) a dynamic and self-correcting statistical approach to combine flu-related Google search frequencies, information from electronic health records, and historical trends within a given state, as well as (2) a data-driven network-based approach that leverages spatial and temporal synchronicities observed in historical flu activity across states to improve state-level flu activity estimates. The proposed ensemble approach considerably outperforms each individual method and any previously proposed state-specific method for flu tracking, with higher correlations and lower prediction errors.

## Introduction

The Internet has enabled near-real time access to multiple sources of medically relevant information, from cloud-based electronic health records to environmental conditions, social media activity, and human mobility patterns. These data streams, combined with an increase in computational power and our ability to process and analyze them, promise to revolutionize the way we treat individual patients and communities in the presence of health threats. As the field of precision medicine [1] continues to yield important medical insights as a consequence of recent improvements in the quality and cost of genetic sequencing as well as advances in bioinformatics methodologies, *precision public health* efforts aim to eventually provide the right intervention to the right population at the right time [2]. In this context, real-time disease surveillance systems capable of delivering early signals of disease activity at the local level may give local decision-makers, such as governments, school districts, and hospitals, valuable and timely information to better mitigate the effects of disease outbreaks. Our work focuses on a methodology aimed at achieving this for influenza activity surveillance.

Influenza has a large seasonal burden across the United States, infecting up to 35 million people and causing between 12000 and 56000 deaths per year [3]. Limiting the spread of outbreaks and reducing morbidity in those already infected are crucial steps for mitigating the impact of influenza. To guide this effort, public health officials, as well as the general public, should have access to localized, real-time indicators of influenza activity. Established influenza reporting systems currently exist over large geographic scales in the United States, coordinated by the Centers for Disease Control and Prevention (CDC). These systems provide weekly reports of influenza statistics, aggregated over the national, regional (10 groups as defined by the Health and Human Services), and starting in fall 2017, state level. Of particular interest, U.S. Outpatient Influenza-like Illness Surveillance Network (ILINet) records the percentage of patients reporting to outpatient clinics with symptoms of influenza-like illness (ILI), which is defined by fever over 100 °F in addition to sore throat or cough, over the total number of patient visits [4]. While these measurements are an established indicator of historical ILI activity, they require around a week to collect from individual health-care providers across the country, analyze, and report and are frequently revised. This delay and potential subsequent revisions can reduce the utility of the system for real-time situational awareness and data analysis.

To address this delay, research teams have devised methods to estimate ILI a week ahead of healthcare-based reports and in near-real time, termed “nowcasting”, at the national and regional levels. These methods incorporate a variety of techniques from statistical modeling and machine learning [5–8], to mechanistic and epidemiological models [8–10]. Many utilize innovative web-based data sources such as Internet search frequencies and electronic health records [5]. Some have also taken into account historically-observed spatial and temporal synchronicities in flu activity [11,12] to improve the accuracy of existing flu surveillance tools [13,14]. Because influenza transmission occurs locally and is spread from person to person, the timing of outbreaks and resulting infection rate curves can significantly differ from state to state. Thus, despite the aforementioned successes in national and regional surveillance, these spatial resolutions are likely not enough to aid decision-making at smaller geographic scales, since important information about localized conditions is lost in regional or national aggregates.

The first influenza nowcasting system at the state level across the United States was Google Flu Trends (GFT). GFT reported a number each week representing influenza activity for each state, various cities, in addition to the national and regional levels, using Google search activity as a predictor. While innovative at the time, studies have pointed out its large prediction errors when tested in real time and proposed alternative methodologies that can incorporate Google searches more effectively at the national level [6,15–17]. A feasibility proposal replacing GFT for flu detection, at the state level, was published last year by Kandula, Hsu, and Shaman, who presented retrospective out-of-sample flu estimates, over the 2005-2011 flu seasons, using a random forest methodology based on Google searches and historical flu activity as predictors [18]. While this study showed promise, the authors did not report significant improvements to GFT and provided only aggregate distributional metrics to evaluate the performance of their models over conglomerates of states (as opposed to state-level metrics), making it challenging to replicate or improve their results for any given state.

### Approach

In this study, we provide a solution for localized flu nowcasting by first extending to each state a proven methodology for inferring flu activity, named ARGO, which combines information from flu-related Google search frequencies, electronic health records, and historical flu trends. Next, we develop a spatial network approach, named Net, which refines ARGO’s flu estimates by incorporating structural spatio-temporal synchronicities observed historically in flu activity. Finally, we introduce ARGONet, a novel ensemble approach that combines estimates from ARGO and Net using a dynamic, out-of-sample learning method. We produce retrospective estimates using ARGO from September 2012 to May 2017 and show that ARGO alone demonstrates strong improvement over GFT and an autoregressive benchmark. Then we generate retrospective flu estimates using ARGONet from September 2014 to May 2017 and show further improvements in accuracy over ARGO in over 75% of the states studied. We present detailed metrics and figures over each state to enable analysis as well as future refinement of our methods.

## Results

### State-level ARGO models outperform existing benchmarks

We first adapted the ARGO (AutoRegression with General Online information) methodology to state-level flu detection. ARGO has previously demonstrated the ability to infer flu activity with high precision over a variety of geographical areas and scales [5,19,20]. The adapted model dynamically fits a regularized multivariable regression on state-level Google search engine frequencies, electronic health record reports from athenahealth, and historical CDC %ILI estimates (see Methods section). We trained a separate ARGO model for each state and used them to generate retrospective out-of-sample estimates from September 30, 2012 to May 14, 2017 for each state in the study.

To assess the predictive performance of ARGO, we also produced retrospective estimates using two benchmarks: a) GFT, a scaled version of the Google Flu Trends time series for each state, fitted to match the scale of each state’s CDC %ILI; and b) AR52, an autoregressive model which uses the CDC %ILI from the previous 52 weeks in a regularized multivariable regression to predict %ILI of the current week (see Methods section for details on both). Since autoregression is an important component of ARGO itself, improvement over AR52 indicates the effective contribution of real-time Google search and electronic health record data. Fig. 1a shows a graphical comparison of ARGO, AR52, and GFT for each state over the time window when GFT estimates were available (September 30, 2012 to August 15, 2015). The three panels display the root mean square error (RMSE), Pearson correlation, and mean absolute percent error (MAPE) over the period (defined under Comparative Analyses in the Methods section). ARGO models outperform GFT in RMSE in every state, in correlation in all but one state, and in MAPE in all but two states. Furthermore, ARGO reduces the RMSE of GFT by > 50% in 23 states and increases correlation by > 10% in 25 states. ARGO also performs comparably or better in RMSE and correlation than AR52, although it does not generally outperform AR52 in MAPE. In all but 8 states, ARGO beats AR52 in a majority (2 or all 3) of metrics (Fig. 1b).

**Fig. 1:**
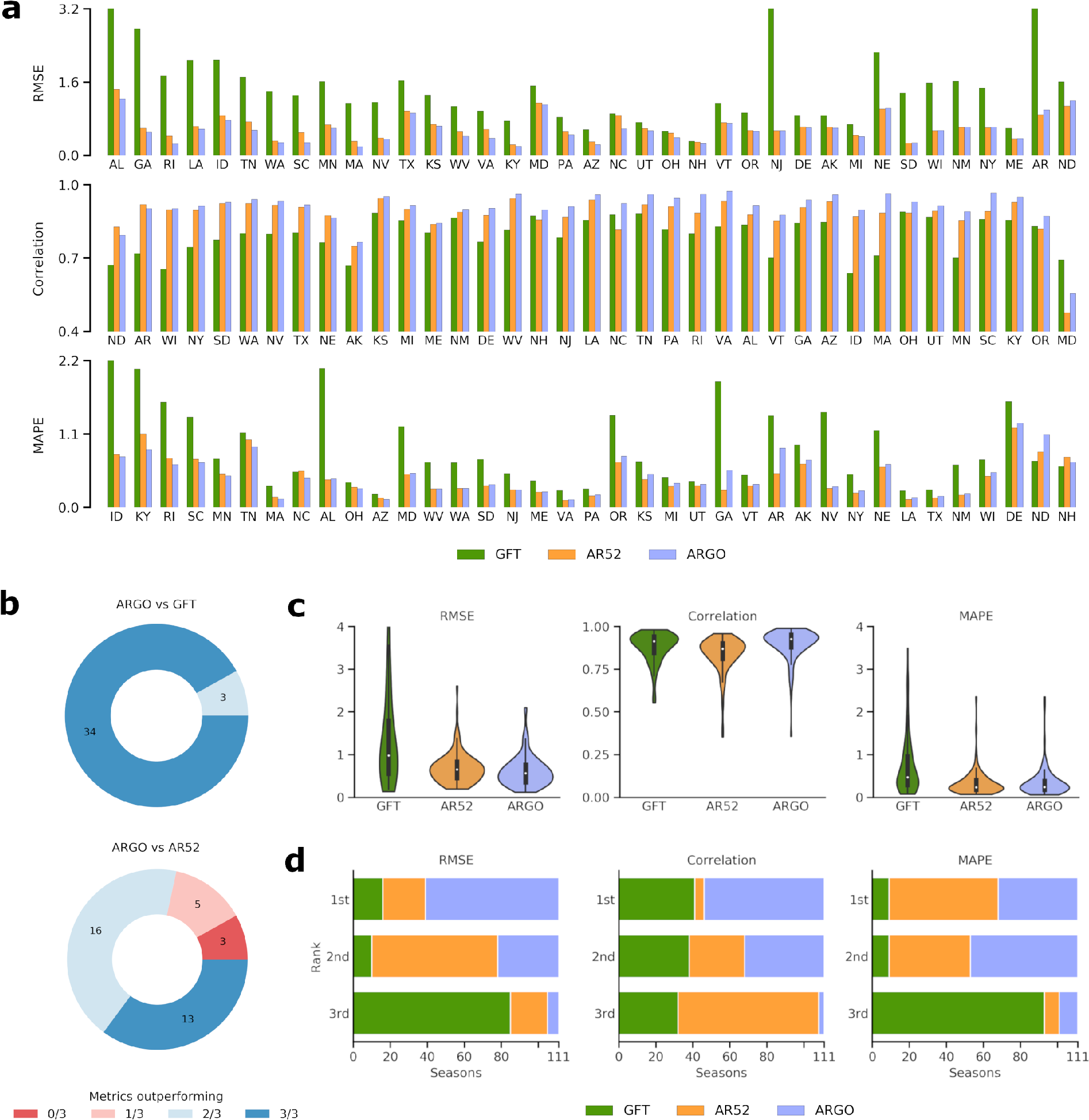
a) State-level performance of ARGO benchmarked with GFT and AR52, as measured by RMSE (top), Pearson correlation (middle), and MAPE (bottom), over the period from September 30, 2012 to August 15, 2015. Extreme GFT error values are displayed up to a cutoff point. States are ordered by ARGO performance relative to the benchmarks to facilitate comparison. b) The proportion of states where ARGO outperforms GFT (top) or AR52 (bottom) in 0, 1, 2, or all 3 metrics. c) The distribution of values for each metric for each model, over the 111 state-seasons during the same period. Numerical values are reported in Table A1. d) The distribution of ranks attained by each model over the 111 state-seasons for each metric.

Attention to flu activity is typically heightened during flu seasons (between week 40 of one year and week 20 of the next), as the majority of seasonal flu cases occur within this time frame. We assessed ARGO performance over each flu season within the study period, namely the 2012-13 to 2014-15 seasons inclusive. With three seasons where comparison with GFT is available and 37 states, this yields 111 state-seasons. Of these, ARGO outperforms GFT in 94 state-seasons in RMSE, 69 in correlation, and 97 in MAPE. ARGO also surpasses AR52 in 83 state-seasons in RMSE, 104 in correlation, and 47 in MAPE (aggregated from Table A3). Correspondingly, ARGO outperforms the benchmarks in terms of median and interquartile range over the seasons, with the exception of MAPE against AR52 (Fig. 1c), and ranks first over the majority of state-seasons in RMSE and correlation (Fig. 1d). Interestingly, despite lower quartile values, GFT has a better tail spread than ARGO in correlation, though not in RMSE or MAPE.

### Incorporating spatio-temporal structure in state-level flu activity

Because ARGO models the flu activity within a given state using only data specific to that state, a natural question is whether information from other states across time can be used to improve the accuracy of flu predictions. As shown in Fig. 2a, historical CDC %ILI observations show synchronous correlations between states. The clustering of intercorrelated states from the same regions suggests that geographical spatio-temporal structure can be exploited as a correctional effect.

**Fig. 2:**
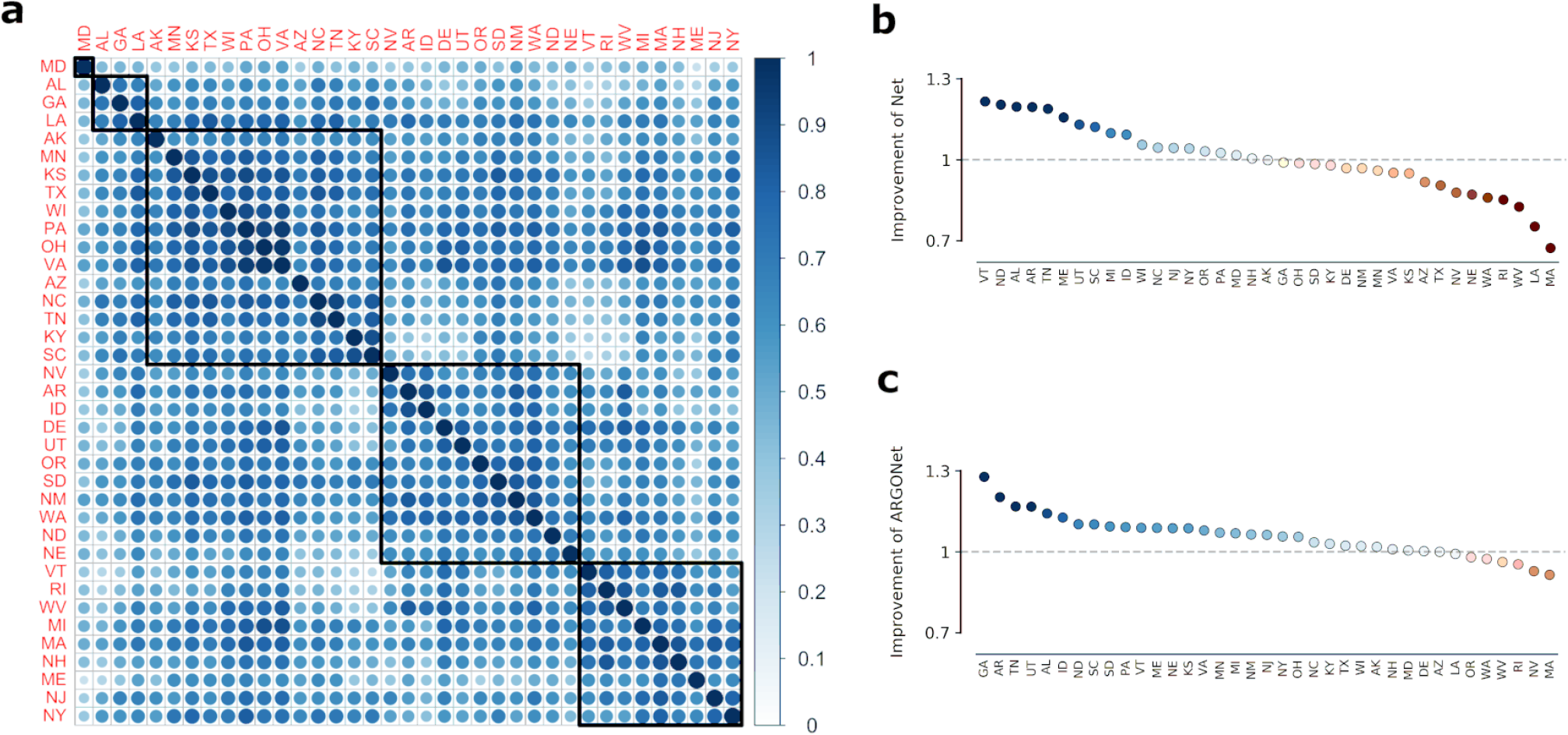
a) Heatmap of pairwise %ILI correlations between all states in the study over the period September 30, 2012 to May 14, 2017. Five clusters of inter-correlated states are denoted by black boxes. b) RMSE improvement of Net over ARGO over the period September 28, 2014 to May 14, 2017. The improvement of Net is here defined as the inverse RMSE ratio of Net and ARGO, so values above 1 indicate improvement. c) RMSE improvement of ARGONet over ARGO over the same period.

Inspired by this finding, we developed a network-based model on each state, which incorporates multiple weeks of historical %ILI activity from all other states in a regularized multivariable regression. Out-of-sample estimates from this model, denoted Net, improve on the RMSE of ARGO on half of the states over the period of Sept. 28, 2014 to May 14, 2017, but show a comparable increase in error on the other half of the states (Fig. 2b). Because ARGO and Net dramatically outperform each other over distinct states, we investigated whether an ensemble combining the relative strengths of each model could lead to significant improvement.

### ARGONet ensemble improves on state-level ARGO models

The resulting ensemble, denoted ARGONet, dynamically selects either ARGO’s or Net’s prediction in each week and state based on the past performance of each model over a suitable training space (see Methods section for details). Over the period where ARGONet estimates were generated (Sept. 28, 2014 to May 14, 2017 after a 2-year training window), we found that this approach resulted in out-of-sample improvement in RMSE over ARGO in all but 8 states (Fig. 2c). Furthermore, in these 8 states, the error increase of ARGONet is relatively controlled compared to the error increase of Net.

In addition to RMSE, ARGONet also displays general improvement in correlation and MAPE over both ARGO and the AR52 benchmark (Fig. 3a). We previously noted that ARGO did not outperform AR52 in MAPE despite being superior in terms of RMSE, which suggests that ARGO is more accurate than AR52 during periods of high flu incidence and less accurate during low flu incidence. On the other hand, by incorporating spatio-temporal structure, ARGONet is able to achieve lower MAPE than AR52 over both the entire time period of Sept. 2014 - May 2017 and the 108 state-seasons within this period (three states are missing %ILI data for the 2016-17 season, resulting in fewer state-seasons compared to the previous analysis) (Fig. 3a-c). Note that while ARGO and Net outperforms the AR52 benchmark by a majority of metrics in 32 and 30 states, respectively, ARGONet does the same in 36 out of 37 states (Fig. 3b).

**Fig. 3:**
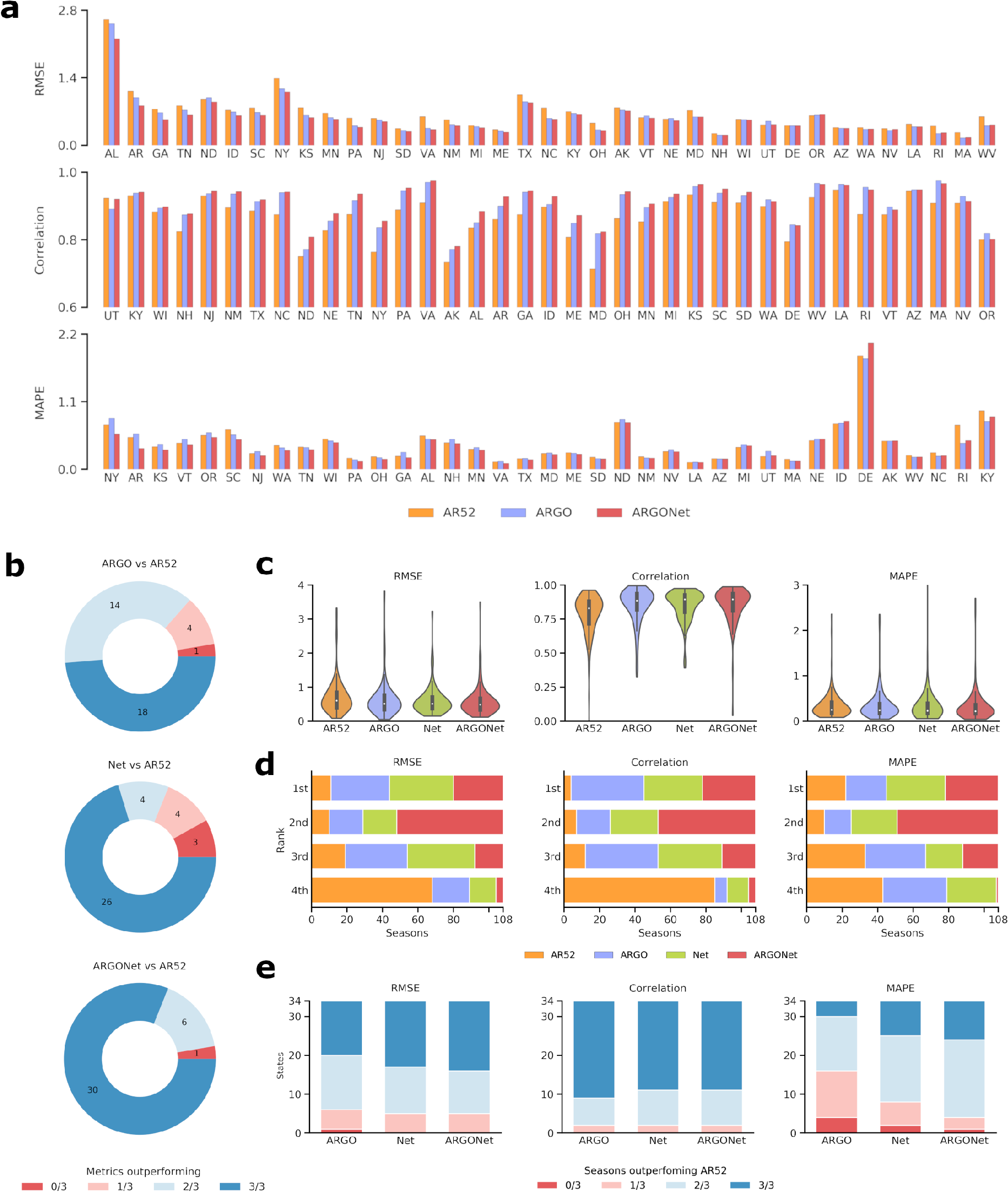
a) State-level performance of ARGONet benchmarked with ARGO and AR52, as measured by RMSE (top), Pearson correlation (middle), and MAPE (bottom), over the period from September 28, 2014 to May 14, 2017. States are ordered by ARGONet performance relative to the benchmarks to facilitate comparison. b) The proportion of states where ARGO (top), Net (middle), or ARGONet (bottom) outperforms AR52 in 0, 1, 2, or all 3 metrics. c) The distribution of values for each metric for each model, over the 108 state-seasons during the same period. Numerical values are reported in Table A1. d) The distribution of ranks attained by each model over the 108 state-seasons for each metric. e) The proportion of states where each model outperforms AR52 in 0, 1, 2, or all 3 flu seasons within the same period. Only 34 states have data for all three flu seasons available in this period.

Interestingly, the performance increase of ARGONet does not appear to stem from being simultaneously more accurate than ARGO and Net over a majority of state-seasons. Note in Fig. 3d that while ARGONet tends to rank first in a smaller proportion of state-seasons than ARGO or Net, ARGONet ranks either first or second in a far larger proportion of state-seasons than the other two models, indicating that the ensemble’s overall success comes from increased consistency. Finally, Fig. 3e subdivides the states by the fraction of seasons (out of 3) where each model outperforms AR52. We see that ARGONet performs favorably (wins 2 or 3 out of 3 seasons) in the vast majority of states, with considerably better distribution in terms of MAPE than ARGO or Net. Refer to Table A3 for numerical metrics over each state and season.

Detailed time series comparisons of ARGO and ARGONet relative to the official CDC-reported %ILI values are shown in Fig. 4. Note that our models consistently track the CDC %ILI curve during both high and low periods of ILI activity, whereas GFT often significantly overpredicts during season peaks. Time series plots specifically comparing ARGONet and ARGO over Sept. 2014 - May 2017 are presented in Fig. A1 and better enable the reader to visually inspect ARGONet’s improvement over ARGO. In concordance with previous results, ARGONet tracks the CDC %ILI curve more accurately than ARGO over some periods of time, while over other periods the curves are identical. This is an expected result of our winner-takes-all ensemble methodology. Some states that especially highlight these patterns are Arkansas (AR), Georgia (GA), New York (NY), and Vermont (VT).

**Fig. 4:**
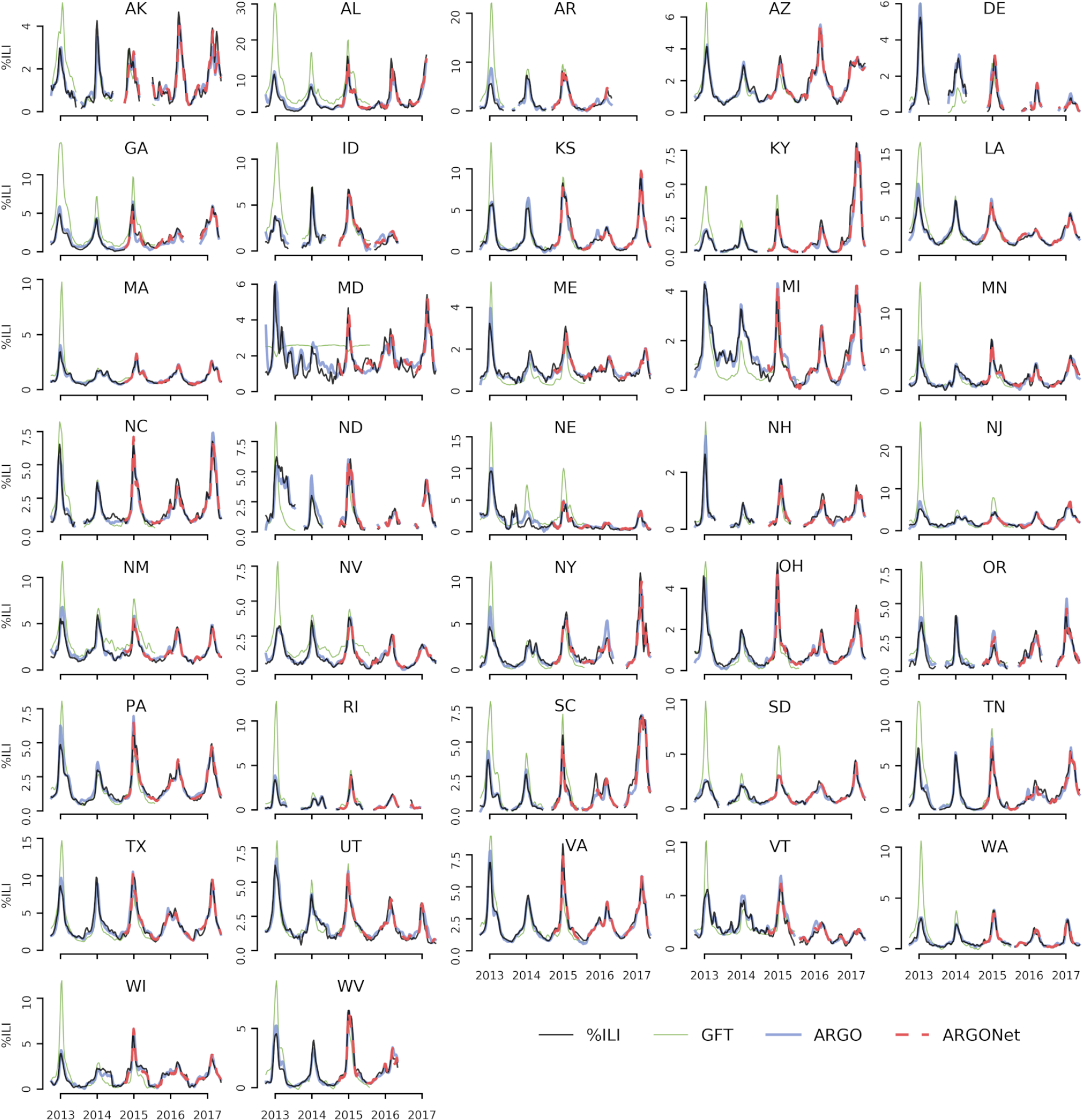
Time series plots displaying the performance of ARGO and ARGONet relative to the official CDC %ILI time series over the entire study period (September 30, 2012 to May 14, 2017). The GFT benchmark is also shown. Refer to Fig. A1 for more detailed figures from Sept. 2014 - May 2017.

## Discussion

Our ensemble, ARGONet, successfully combines Google search frequencies and electronic health record data with spatial and temporal trends in flu activity to produce accurate forecasts for current ILI activity at the state level. We believe that the accuracy of our method involves a balance between responsiveness and robustness. Real-time data sources such as Google searches and electronic health records provide information about the present, allowing the model to immediately respond to current flu trends. On the other hand, using the values of past CDC flu reports in an autoregression adds robustness by preventing our models from creating outsize errors in prediction. Similarly, incorporating spatial synchronicities adds stability by maintaining state-level inter-correlations evident in historical flu activity. Our results suggest that dynamic learning ensembles incorporating real-time Internet-based data sources can surpass any individual methods in inferring flu activity.

Previous work has shown the versatility of ARGO, one of the component models in our ensemble, over a variety of disease estimation scenarios [5,20–22]. At the state level, where it had not been applied before, it clearly outperformed existing benchmarks over the study period, namely Google Flu Trends and an autoregression. While ARGO alone performs better than study benchmarks, we also found that spatio-temporal synchrony could be used to improve model accuracy (Net). Combining web-based data sources with this structural network-based approach (ARGONet) further improves prediction accuracy and suggests the future study of synchronous network effects at varying geographical scales. Future work should explore adding similar approaches to flu nowcasting systems at finer spatial resolutions, such as the city level.

Accurate flu monitoring at the state level faces challenges due to higher variance in data quality across states. The official %ILI reporting system within each state varies in reporting coverage and consistency, and thus the magnitudes of flu activity may not reflect actual differences of flu activity between states. In addition, the quality of Google Trends frequencies and the prevalence of clinics reporting to athenahealth (the provider of our electronic health record data) vary considerably from state to state, affecting the ability of our models to extract useful information from these data sources. In past work, we hypothesized that better model performance could be explained as a consequence of higher Internet-based data quality. As a result, we believe that geographical improvement of ARGONet over the benchmark AR52 (as defined by percent reduction of RMSE) may also be associated with factors that serve as proxies of Google Trends or athenahealth data quality, namely detectable flu-related search terms from Google Trends and estimated athenahealth population coverage (Fig. A2a-d). Regression analysis indicates a moderate association of ARGONet improvement with athenahealth coverage (Fig. A2e) and a weak association with detectable flu-related Google search activity (Fig. A2f). Interestingly, flu-related search activity has a very strong correlation with state population (Fig. A2g), which suggests that larger pools of Internet users result in better signal-to-noise ratio in search activity. Future analysis can examine the interplay of these factors with CDC %ILI report quality and structural spatial correlations. For example, we hypothesize that the Net model contributes strongly in states with lesser-quality data which are adjacent to states with high-quality data. Clustering states by pairwise %ILI correlation (Fig. 2a) shows that geographic proximity is a relevant synchronous factor. Within the boxed clusters, we see that Southeastern states, Western states, and New England/mid-Atlantic states often group together.

Localized, accurate surveillance of flu activity can set a foundation for precision public health in infectious diseases. Important developments in this field can involve emerging methodologies for tracking disease at fine-grained spatial resolutions, rapid analysis and response to changing dynamics, and targeted, granular interventions in disparate populations, each of which has the potential to complement traditional public health methods to increase effectiveness of outcomes [23]. We believe that the use of our system can produce valuable real-time subregional information and is a step toward this direction. At the same time, the performance of ARGONet depends directly on the availability and quality of Internet-based input data and also relies on a consistently reporting (even if lagged) healthcare-based surveillance system. We anticipate that data sources will improve over time, for example, if athenahealth continues to gain a larger market share over the states or more Google Trends information becomes available. If these conditions hold, or as more web-based data sources including other electronic health record systems become available in real time, the accuracy of our methods may continue to increase.

## Methods

### Data Acquisition

Three data sources were used in our models: influenza-like illness rates from ILINet, Internet search frequencies from Google Trends, and electronic health records from athenahealth. Weekly information from each data source was collected for the time period of October 4, 2009 to May 14, 2017.

### Influenza-like illness rates

Weekly influenza rates reported by outpatient clinics and health providers for each available state were used as the epidemiological target variable of this study. The weekly rate, denoted %ILI, is computed as the number of visits for influenza-like illness divided by the total number of visits. Data from October 4, 2009 to May 14, 2017 for 37 states were obtained from the CDC. For inclusion in the study, a state must have data from October 2009 to May 2016, with no influenza seasons (week 40 of one year to week 20 of the next) missing. Some states were missing data, usually due to not reporting in the off-season (between week 20 and week 40 of each year). Missing or unreported weeks, as well as weeks where 0 cases were reported, were excluded from analysis on a state-by-state basis. While in real-time ILI values for a given week may be revised in subsequent weeks, we only had access to the revised version of these historical values.

### Internet search frequencies

Search volumes for specific queries in each state were downloaded through Google Trends, which returns values in the form of frequencies scaled by an unknown constant. While our pipeline used the Google Trends API for efficiency, search volumes can be publicly obtained from www.trends.google.com for reproducibility. Relevant search terms were identified by downloading a complete set of 287 flu-related search queries for each state, and keeping the terms that were not completely sparse. Because Google Trends left-censors data below an unknown threshold, replacing values with 0, a query with high sparsity indicates low frequency of searches for that query within the state.

In an ideal situation, relevant search queries at the state-level resolution would be obtained by passing the historical %ILI time series for each state into Google Correlate, which returns the most highly correlated search frequencies to an input time series. However, such functionality is only supported at the national level, at least in the publicly accessible tool. Given this limitation, we used two strategies to select search terms:

1. A initial set of 128 search terms was taken from previous studies tracking influenza at the US national level [6].
2. To search for additional terms, we submitted multiple state %ILI time series into the Google Correlate and extracted flu-related terms, under the assumption that some of the state-level terms would show up at the national level.

To minimize overfitting on recent information, the %ILI time series inputted into Google Correlate were restricted from 2009-2013. State-level search frequencies for the union of these terms and the 128 previous terms were then downloaded from the Google Trends API, resulting in 282 terms in total (Table A2).

### Electronic health records

Influenza rates for patients within the athenahealth provider network are provided weekly from athenahealth on each Monday. Three types of syndromic reports were used as variables: ‘influenza visit counts’, ‘ILI visit counts’, and ‘unspecified viral or ILI visit counts’, which were converted into percentages by dividing by the total patient visit counts for each week. The athenahealth network and influenza rate variables are detailed in Santillana et al. [19]

### Google Flu Trends

In addition to the above data sources, we downloaded GFT estimates as a benchmark for our models. GFT provided a public flu prediction system for each state until its discontinuation in August 2015 [24]. GFT values were downloaded and scaled using the same initial training period of 104 weeks used in all of our models (October 4, 2009 to September 23, 2012).

## Models

### ARGO

The time series prediction framework ARGO (AutoRegression with General Online information) issues flu predictions by fitting a multivariable linear regression each week on the most recent available Internet predictors and the previous 52 %ILI values. Because of many potentially redundant variables, L1 regularization (Lasso) was applied to produce a parsimonious model by setting the coefficients for weak predictors to 0. The model was re-trained each week on a shifting 104-week training window in order to adapt to the most recent two years of data, and the regularization hyperparameter was selected using 10-fold cross validation on each training set. Details about the ARGO model and its applicability in monitoring infectious diseases such as influenza, dengue, and zika are presented in previous work [5,21,22]. Refer to the Appendix for a detailed mathematical formulation of ARGO.

To fine-tune predictive performance, adjustments to ARGO were introduced on a state-by-state basis:

- Filtering features by correlation: For each week, non-autoregressive features ranked outside the top 10 by correlation were removed to reduce noise from poor predictors. Based on previous research, this complementary feature selection process benefits the performance of lasso, which can be unstable in variable selection [20, 21].
- Regularization hyperparameters: Features with high correlation to the target variable over the training set received a lower regularization weight, which makes them less subject to the L1 penalty (see the Appendix for derivation).
- Weighting recent observations: Although ARGO dynamically trains on the last 104 weeks of observations, more recent observations likely contain more relevant information. Thus the most recent 4 weeks of data received a higher weight (set to be twice the weight of the other variables) in the training set.

### Network-based approach

Historical CDC ILI observations show synchronous relationships between states, as shown in Fig. 2. To identify these relationships with the goal of improving our %ILI predictions, for each state, we dynamically constructed a regularized multi-linear model for each week that has the following predictors: %ILI terms for the previous 52 weeks for the target state, and the synchronous (same week’s values) and the past three week’s of observed CDC’s %ILI terms for the other states. Notice that to produce predictions of %ILI for a given state in a given week, the model requires synchronous %ILI from the other states, which would not be available in real-time. Instead, we used ARGO predictions for the current week as surrogates for these unobserved values. This model was re-trained weekly using all previously observed data with 10-fold cross-validation to determine the L1 regularization term (formulation in Appendix). This model is denoted Net.

### Ensemble approach

In order to optimally combine the predictive power of ARGO and Net, we trained an ensemble approach based on a winner-takes-all voting system, which we named ARGONet. ARGONet’s prediction for a given week is assigned to be Net’s prediction if Net produces lower error relative to the observed CDC %ILI (specifically root mean square error, as defined in Comparative analyses) in the previous *K* predictions as compared to ARGO. Otherwise, ARGONet’s prediction is assigned to be ARGO’s prediction. To determine the hyper-parameter *K* for each state, we trained ARGONet using the first 104 out-of-sample predictions of ARGO. Here *K* can take the value of either 1, 2 or 3. The value of *K* that yielded the lowest RMSE between ARGONet and CDC’s %ILI over the training period was chosen to produce out-of-sample predictions in the unseen time period (September 28, 2014 onward).

### Comparative analyses

To assess the predictive performance of the models, we produced state-level retrospective estimates using two benchmarks: a) “AR52”, an autoregressive model, which uses the %ILI from the previous 52 weeks in a LASSO regression to predict %ILI of the current week, and b) “GFT”, made by scaling each state’s Google Flu Trends time series to its official revised %ILI using the same initial training period of 104 weeks used in our models as discussed in the Methods section.

The performances of all models and benchmarks compared to the official (revised) %ILI were scored using three metrics: root mean squared error (RMSE), Pearson correlation coefficient, and mean absolute percent error (MAPE). These were computed over the entire study period (September 30, 2012 to May 14, 2017) and over each influenza season (defined as week 40 of one year to week 20 of the next) within the study period.

The models and benchmarks were further scored over two specific sub-periods: 1) the window when GFT was available (September 30, 2012 to August 9, 2015), and 2) the window starting with the first available ARGONet prediction (September 28, 2014 to May 14, 2017).

ARGO model estimates were generated using scikit-learn in Python 2.7 [25], while Net and ensemble models were generated in R 3.4.1. Analysis was conducted in Python except for Fig. 2a, which used the R package corrplot [26].

## Acknowledgments

This work was partially funded by the Centers for Disease Control and Prevention’s Cooperative Agreement PPHF 11797-998G-15. The authors thank Josh Gray, Anna Zink, and Dorrie Raymond for the collection and processing of the data from athenahealth.

The authors would also like to thank Matthew Biggerstaff from the National Center for Immunization and Respiratory Disease, Centers for Disease Control and Prevention, for great insights and guidance on the direction and objectives of this study.

## Appendix ARGO formulation

Let *y*_*i,t*_ be the CDC %ILI in state *i* at time *t*, and let ***X***_*i,t*_ = {*X*_*i,t,k*_}_*k*∈{1,…,*M*}_ be the vector of Internet-based data in corresponding state and time. ARGO assumes a hidden Markov model based on an autoregressive structure with *N* lags, as shown below:

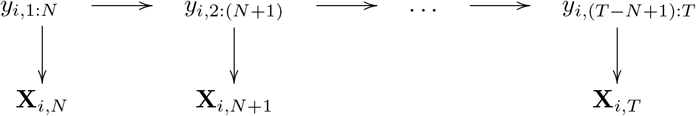

Here, the vectors {*y*_*i*_,_(*t*−*N* +*1*):*t*_}_*t*≥*N*_ follow the Markov property, and at time *T*, the *T*th such vector has not yet been fully observed. Meanwhile, the observed variables **X**_*i,T*_ depend only the corresponding hidden *y*_*i,T*_.

## ARGO model

The above formulation results in the model

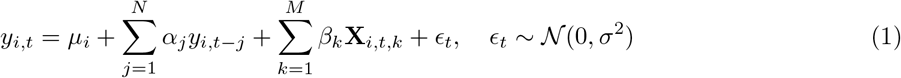

We take *N* = 52 to incorporate short-term and seasonal autoregressive trends within the past year of data, and *M* = 285 at maximum, corresponding to the number of Google Trends and athenahealth variables. The high number of input variables gives us a *p* > *n* situation, so we impose *L*_1_ regularization. Therefore, we can solve for parameters *μ*_*i*_, *α* = (*α*_1_,…,*α*_*N*_), and *β* = (*β*_1_,…,*β*_*M*_) which minimize the objective function

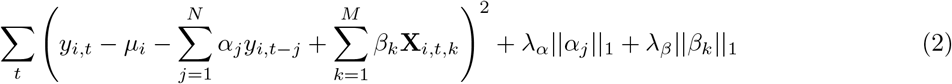

using a rolling training window consisting of the 104 weeks prior to time *t*, with hyperparameters λ_*α*_ and λ_*β*_.

## Hyperparameters

The parameters in (2) are governed by hyperparameters λ_*α*_ and λ_*β*_; however, we introduced a modification to the model. Rather than adhering to the groups *α* and *β*, we replaced them with more flexible groups in the following manner:

Let γ = {*α*_1_,…,*α*_N_,*β*_1_,…,*β*_*M*_} be the set of regularized parameters. Take *p* ⊆ γ to be a set of “priority” parameters, and *q* = γ − *P* to be the remaining parameters. Letting γ_P_ and γ_q_ represent the parameter vectors corresponding to these sets, the objective function simply becomes

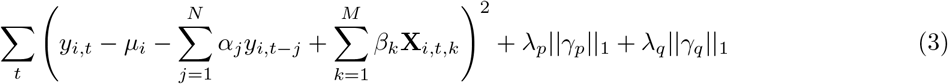

To limit the model space, we allowed *p* to take one of 5 configurations and selected the one with best out-of-sample performance within each state. The configurations are the parameters corresponding to:

1. the 10 highest correlated input variables to the %ILI vector over the training set
2. athenahealth variables
3. athenahealth variables and the most correlated autoregressive terms, namely {*y*_*t−i*_}_*i*∈{1,2,3,4,12,26,52}_
4. the 3 most correlated Google Trends variables over the training set and the most correlated autoregressive terms
5. athenahealth variables, the 3 most correlated Google Trends variables, and the most correlated autoregressive terms.

To further constrain the search space, we maintain a hyperparameter ratio λ_*p*_/λ_*q*_ = 1/10. Since λ_*p*_ is smaller, this allows the priority parameters to take larger values, effectively giving them more weight in the regression. The single hyperparameter was then determined using 10-fold cross-validation over the training set. In practice, prediction accuracy was robust to the specific value of the ratio.

## Net formulation

While the ARGO model for any given state *i* is constrained to the data within state *i*, we hypothesized that *y*_*i,t*_ shows spatio-temporal structure with other states *s* ≠ *i* in the short term (i.e. over the past 4 weeks). Adopting the previous notation, the Net model is then

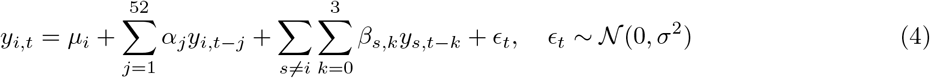

This model was fit using an expanding training window consisting of all previously observed data with 10-fold cross-validation for the regularization hyperparameter. However, for prediction at a given time *t*, the concurrent %ILI values *y*_*s,t*_ are not yet observed, so we replace them with the corresponding real-time ARGO estimates 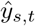. This substitution inherently assumes that ARGO is unbiased, i.e. 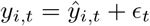, 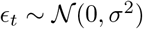.

**Fig. A1:**
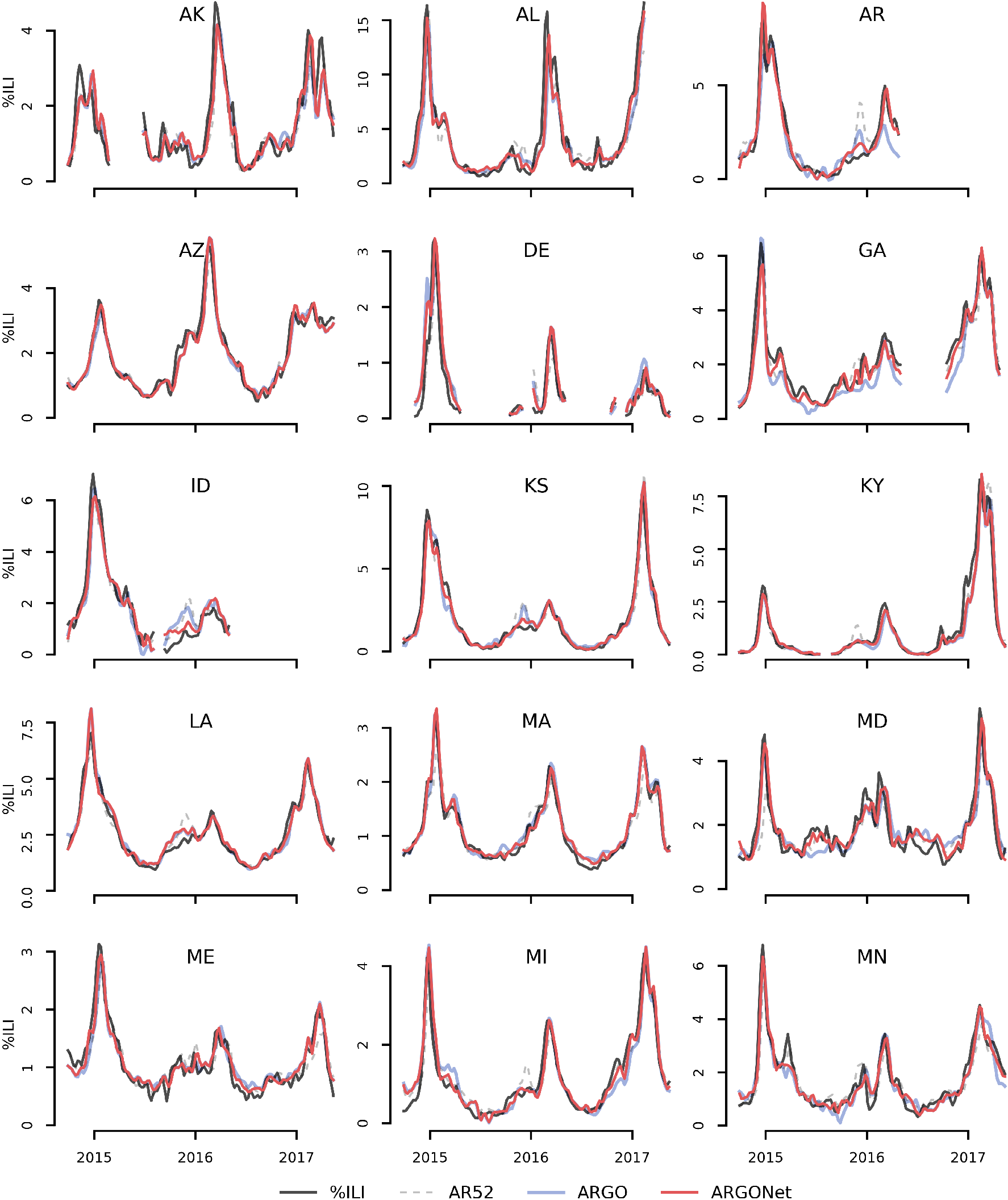
Time series plots displaying the performance of ARGO and ARGONet relative to the official CDC %ILI time series over the period September 28, 2014 to May 14, 2017. The AR52 benchmark is also displayed.

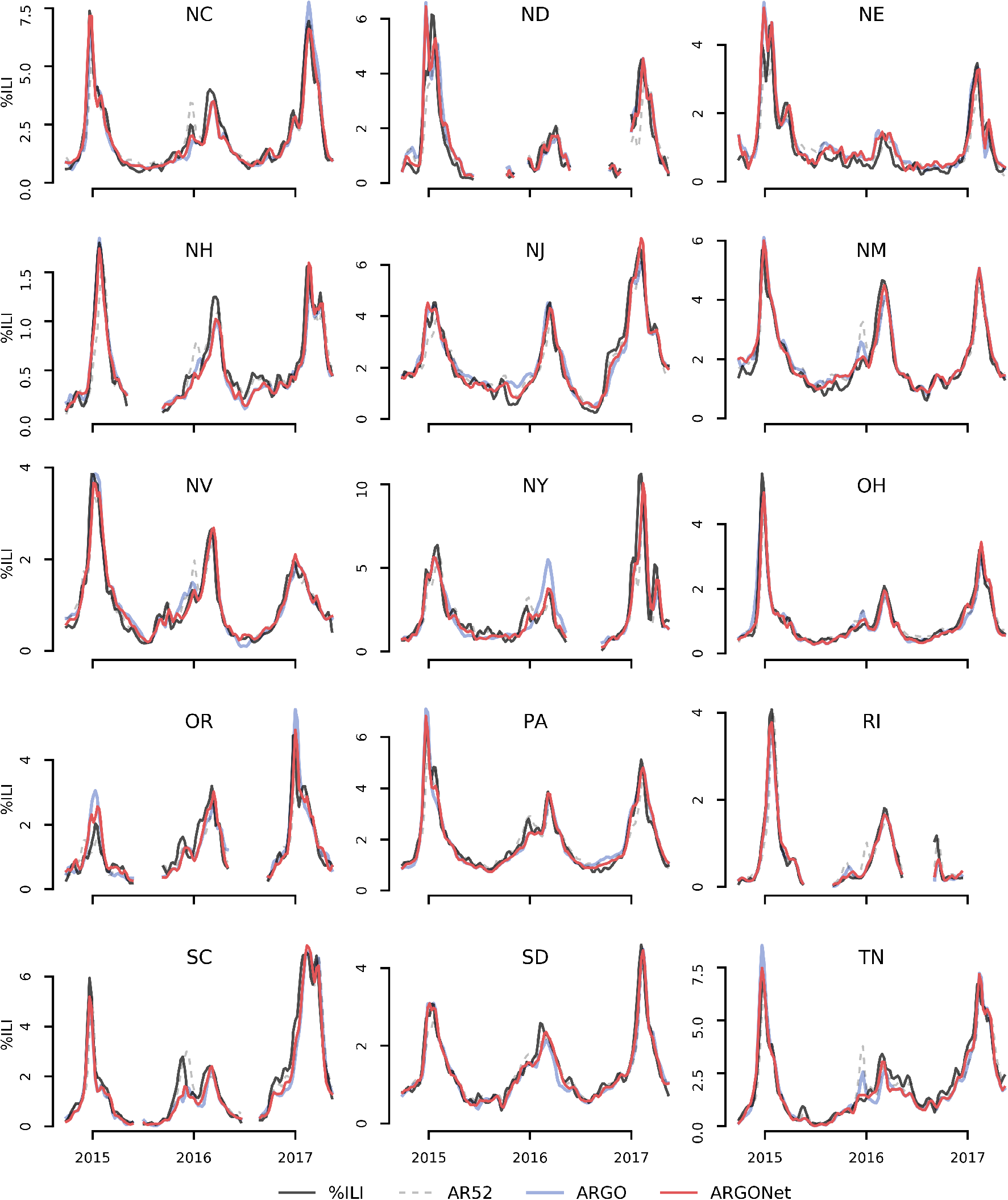

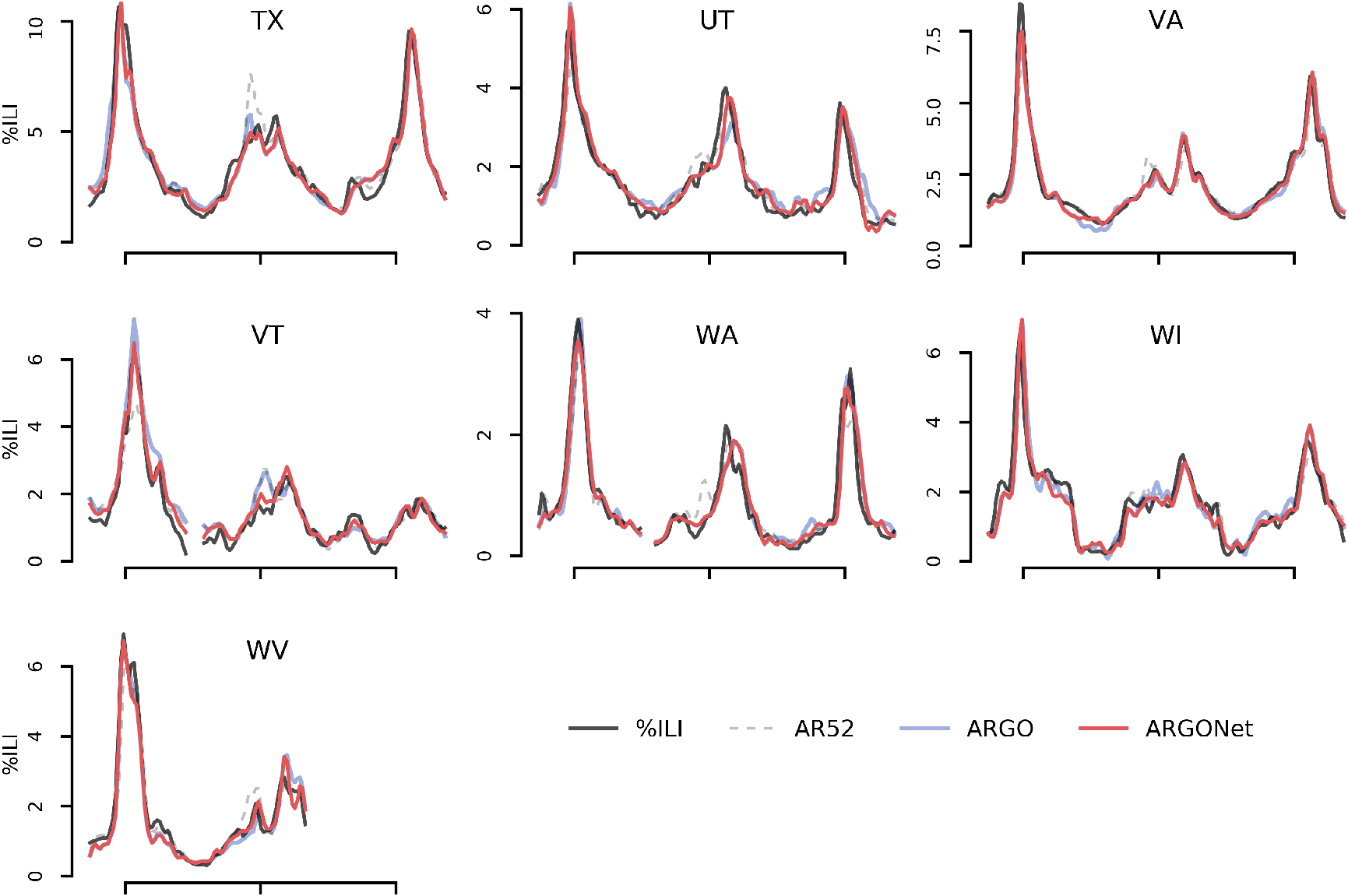

**Fig. A2:**
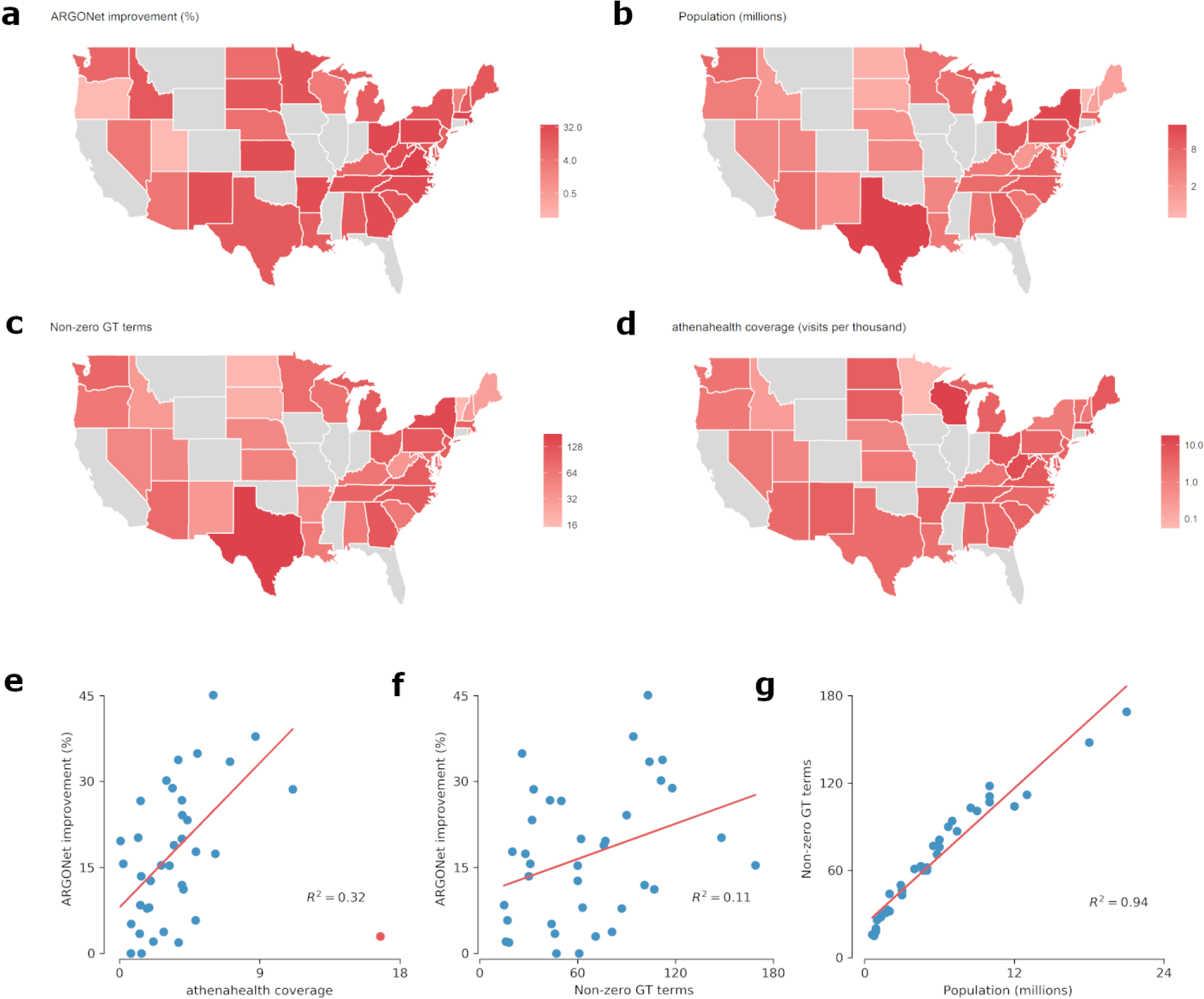
a) Geographical heatmap of the improvement (%RMSE reduction) of ARGONet over AR52. Possible explanatory factors for improvement over an autoregressive benchmark are b) population, which can affect data quality or spatial structure, c) the number of detectable flu-related Google Trends terms per state, which can be taken as a proxy of search data quality, and d) athenahealth coverage per state, calculated as average visits per thousand. e-g) Regression analysis indicates the presence of associations between e) ARGONet improvement and athenahealth coverage, with one outlier (Wisconsin, colored red) removed, f) ARGONet improvement and Google Trends quality, and g) state population and Google Trends quality.

**Table A1:** Aggregate metrics over each flu season for each state corresponding to the violin plots in Figures 1c and 3c. The reported numbers are the median and interquartile ranges of the aggregates. The final two rows are the best reported aggregates for each metric from Kandula et al. for comparison. These were not discussed in the main paper because of the different time period.

| Model | RMSE | Correlation | MAPE |
| --- | --- | --- | --- |
| 2014-15 to 2016-17 seasons |  |  |  |
| ARGONet | 0.48 (0.33-0.67) | 0.90 (0.81-0.94) | 0.22 (0.12-0.36) |
| Net | 0.51 (0.37-0.71) | 0.90 (0.80-0.93) | 0.23 (0.17-0.39) |
| ARGO | 0.51 (0.34-0.75) | 0.89 (0.82-0.94) | 0.24 (0.18-0.38) |
| AR52 | 0.60 (0.39-0.85) | 0.83 (0.72-0.88) | 0.25 (0.18-0.42) |
| 2012-13 to 2014-15 seasons |  |  |  |
| ARGO | 0.56 (0.36-0.77) | 0.93 (0.88-0.95) | 0.25 (0.18-0.38) |
| AR52 | 0.65 (0.46-0.84) | 0.87 (0.81-0.90) | 0.24 (0.18-0.40) |
| GFT | 0.98 (0.55-1.80) | 0.91 (0.84-0.94) | 0.47 (0.29-0.96) |
| 2005-06 to 2010-11 seasons |  |  |  |
| GFT | 0.93 (0.66-1.33) | 0.89 (0.80-0.94) | 0.71 (0.44-1.51) |
| Kandula et al. | 0.84 (0.54-1.25) | 0.86 (0.75-0.91) | 0.54 (0.33-0.90) |

**Table A2:**
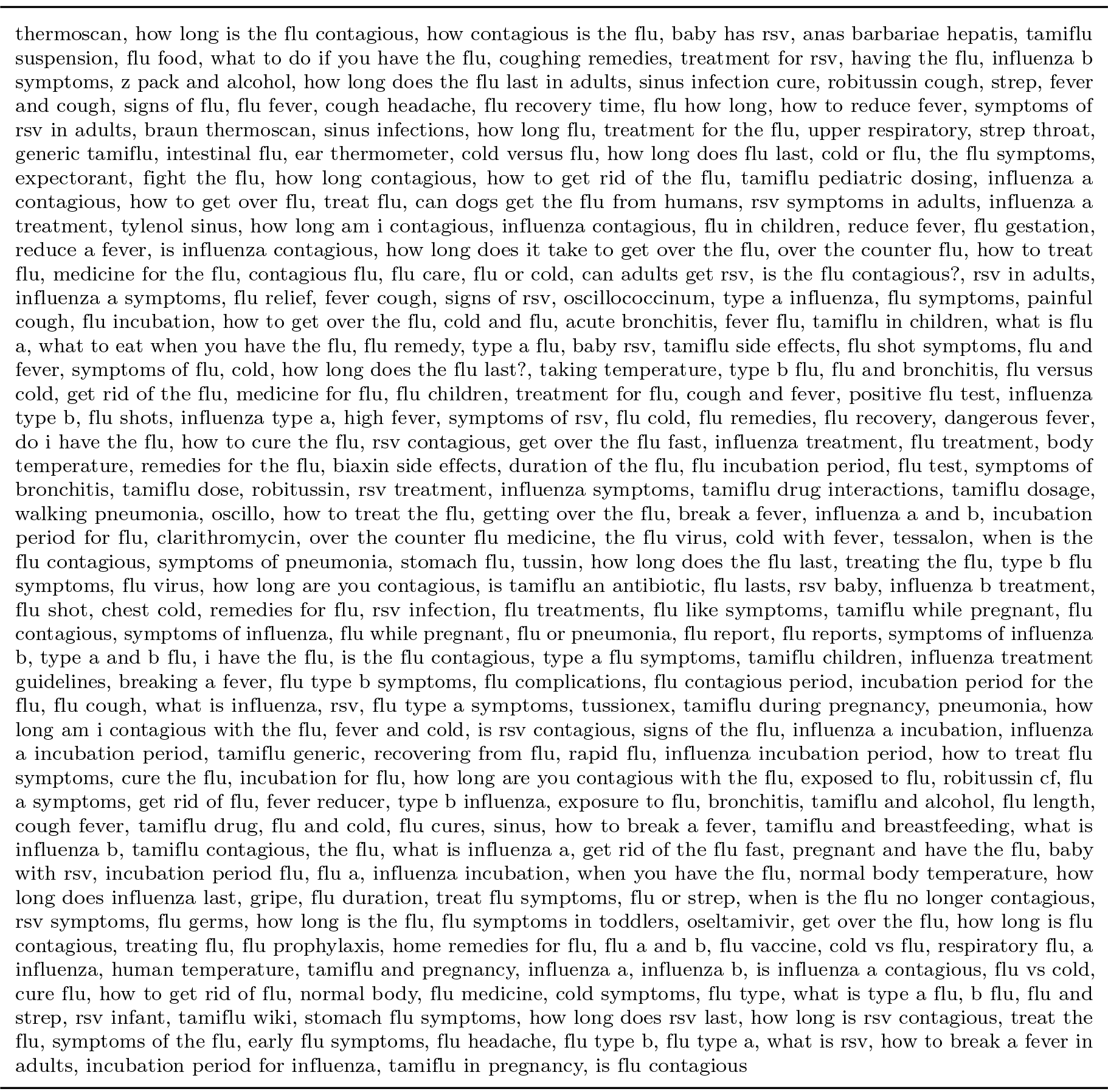
Complete search queries downloaded from Google Trends

**Table A3:** See separate supplementary excel file.

